# 3D genome remodelling underlies PU.1-dependent granulocyte maturation

**DOI:** 10.64898/2026.07.21.739714

**Authors:** Wendy Jia, Timothy M. Johanson, Aleksandar Dakic, Alexandra L. Garnham, Gordon K. Smyth, Stephen L. Nutt, Rhys S. Allan, Hannah D. Coughlan

## Abstract

PU.1 is an ETS-domain transcription factor that has critical roles in many aspects of hematopoiesis and immune cell fate and function. In addition, aberrant PU.1 expression has been implicated in the development of acute myeloid leukemia (AML). Loss of PU.1 during adult murine hematopoiesis results in the expansion of immature granulocytes suggesting that PU.1 plays an important role in granulocyte maturation. To understand the molecular underpinnings of this process, we combined gene expression, transcription factor binding and 3D genome analyses with conditional deletion of PU.1 *in vivo.* We find that in contrast to normal granulocytes, PU.1-deficient cells possessed a transcriptome of immature granulocytes, in line with their cellular phenotype. Furthermore PU.1-deficient granulocytes display altered 3D genome architecture with a significant loss of interactivity in regions bound by PU.1 in control cells. Overall, this study implicates PU.1 as a key regulator of granulocyte maturation and lineage commitment through control of transcriptional programs and 3D chromatin architecture.

## Introduction

Purine-rich box 1 (PU.1) is an ETS-family transcription factor encoded by the *Spi1* gene that has been implicated in many aspects of immune system development and function. Numerous studies have reported that genetic ablation of PU.1 results in profound alterations in hematopoiesis due to a loss of lymphoid and myeloid progenitor populations^1, 2, 3^. This is thought to be largely due to PU.1 directly regulating developmentally important cytokine receptors such as M-CSFR, GM-CSFR and IL7Rα^4^. A consequence of PU.1 inactivation is also a developmental block in granulopoiesis leading to an accumulation of immature granulocytes^1, 5, 6^. Such a buildup may establish a preleukemic state as the loss of PU.1 leads to the development of acute myeloid leukemia (AML) with the appearance of immature granulocytes^7, 8^. In humans, while PU.1 is infrequently mutated in AML, modest changes in the levels of PU.1 has been linked to the development of a preleukemic state^9^. Interestingly, recent studies demonstrated that reintroduction or pharmacological redistribution of PU.1 can promote AML differentiation to overcome this block leading to tumor cell death^10, 11^.

PU.1 is a pioneer transcription factor. It can bind its DNA motif in a repressive chromatin context, leading to an ability to remodel chromatin^12^. As such, it is generally associated with positively regulating gene expression in combination with other transcription factors such as RUNX, C/EBP and AP1 family members to influence lineage specification and immune reactions. Given its pioneering function, ectopic expression of PU.1 in combination with C/EBPα enables reprogramming of fibroblasts, T or B cells to macrophages^13^. While PU.1 often directly binds to promoters of its target genes, it is known to have many binding sites outside of promoters such as in distal enhancer elements^14, 15, 16^. Several groups have provided evidence that PU.1 binding can promote chromatin accessibility of enhancers leading to the deposition of active or permissive histone modifications^14, 15, 17^. Enhancers frequently form long-range interactions or loops with their target gene promoter to regulate transcription^18^, raising the possibility that PU.1 may be involved in establishing the 3D genome to guide the immune cell fate and function. In support of this, PU.1 and other pioneer transcription factors have been shown to cooperate with CCCT-binding factor (CTCF)^19^, which plays a critical role in the formation of chromatin structure. Indeed, PU.1 has been implicated in chromosome looping of specific gene loci^20, 21^. Moreover, a recent study on mature human neutrophils used naturally occurring single nucleotide polymorphisms affecting the DNA binding motifs of PU.1 to suggest that PU.1 has a role in establishing or maintaining gene regulatory enhancer-promoter interactions^22^. However, how the 3D genome is affected at a global scale by the loss of PU.1 during a developmental process such as hematopoiesis has yet to be formally examined. Here we conditionally inactivated PU.1 *in vivo* to study its role in 3D genome conformation during granulopoiesis. We found that PU.1-deficient granulocytes did not establish the 3D genome architecture and corresponding transcriptional program of mature control granulocytes. We observed that chromatin interactions that were associated with PU.1 binding in control granulocytes were not established in the absence of PU.1. Overall, this data implicates PU.1 in directing granulocyte maturation via the establishment of the appropriate 3D genome and transcriptional programs.

## Results

### In vivo PU.1 inactivation results in aberrant granulocyte maturation at the cellular and transcriptional level

To identify the molecular changes that result from the inactivation of PU.1 during granulopoiesis we crossed mice with LoxP sites flanking exon 5 of the *Spi1* gene to tamoxifen-inducible CreER^T2^ mice^1, 23^. Seven and 14 days after tamoxifen treatment we purified granulocytes from the bone marrow of these mice using fluorescently activated cell sorting (Fig. 1A). As expected, control and PU.1-deficient granulocytes both expressed Ly6G and Ly6C, however, PU.1-deficient granulocytes lacked the expression of PU.1-target gene product CD11b (Mac1) and maintained the expression of cKit (Fig. S1A). We confirmed that indeed these cells had minimal expression of PU.1 using qPCR (Fig. S1B).

**Fig. 1.**
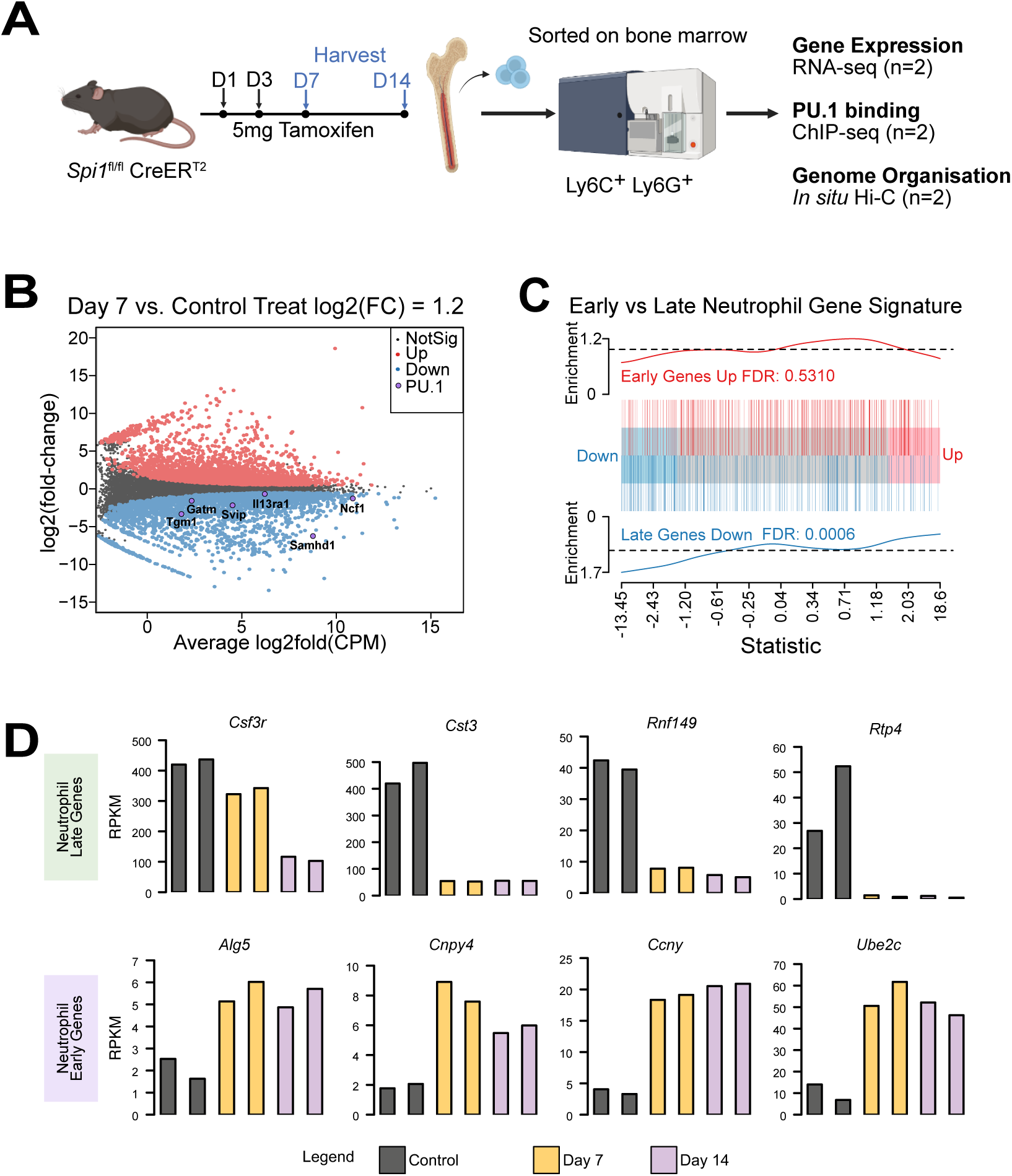
Transcriptome wide alterations in PU.1 deficient granulocytes. **A**, Experimental schematic showing granulocytes were harvested from *Spi1*^fl/fl^ CreER^T2^ mice following PU.1 deletion at day 7 and 14 post-tamoxifen treatment. Controls were untreated *Spi1*^fl/fl^ CreER^T2^ mice. Replicates (n=2 per group) were performed for molecular assay. **B**, Mean-Difference (MD) plot showcasing log2FC comparison between control and day 7 samples against mean abundance (log2 counts per million, CPM). Significantly differentially expressed genes (P-value < 0.05) are colour coded. Candidate PU.1 target genes were also highlighted in the plot. **C**, Barcode plots represent the enrichment of early/late neutrophil gene signatures. Each straight bar corresponds to a specific gene (red for neutrophil early genes and blue for late neutrophil gene) and the line above correspond to relative enrichment. Statistical testing was performed using the fry method described in limma. **D**, Bar plots representing expression profiles of genes as reads per kilo-base (RPKM) with biological replicate samples shown.

RNAseq analysis revealed ∼6300 differentially expressed (DE) genes in PU.1-deficient granulocytes 7 days after tamoxifen treatment versus control (Fig. 1B) with 3128 up regulated genes and 3166 down regulated genes. Remarkably these alterations were stable with very few DEs present between days 7 and 14 (Fig. S1C), suggesting a phenotypic stabilisation had occurred. The genes we found to be deregulated were classic PU.1 target genes including *Gapt*, *Svip*, *Nfam1*, *Samhd1*^24^ (Fig. 1B). Previously we have demonstrated that conditional PU.1 deletion leads to the accumulation of immature granulocytes resembling myeloblasts^1, 7^, therefore we sought to understand if this could be observed at the level of gene expression. We compared our granulocytes to publicly available RNAseq profiles of early and late stages of neutrophil development (Fig. 1C). We observed that PU.1-deficient granulocytes displayed significantly reduced expression of late-stage neutrophil signature genes (Fig. 1C, FDR 0.0006), such as *Csf3r*, *Cst3*, *Rnf149* and *Rtp4* (Fig. 1D) and a trend towards increased expression of early-stage neutrophil genes (Fig. 1C), such as *Alg5*, *Cnpy4*, *Ccny*, *Ube2c* (Fig. 1D). Overall, we conclude that PU.1 is required for the gene expression underlying granulocyte maturation and that *in vivo* deletion of PU.1 in adult progenitors results in the accumulation of cells that display transcriptional hallmarks of granulocytes that have not reached maturity.

### PU.1 binding is predominantly associated with active genes in mature granulocytes

We next sought to examine how PU.1 is involved in regulating gene expression during granulocyte maturation by examining its genomic localisation using chromatin-immunoprecipitation sequencing (ChIPseq). We found that PU.1 was bound to 9001 sites in the wildtype granulocyte genome, with 3932 (44%) of these being at promoters (Fig. 2A and Fig. S2A). We observed a significant correlation between PU.1 binding in promoters and downregulation in PU.1-deficient granulocytes (Fig2B, Fig. S2B, Fry test p-value). Further analysis showed that of the ∼2000 DE genes that had PU.1 promoter binding >70% were downregulated after PU.1 inactivation compared to 45% of unbound DE genes (Fig. S2C, p-value from chi-sq test). Overall, the expression of PU.1-bound genes were significantly reduced in log2FC in the PU.1-inactivated granulocytes as compared to unbound genes (Fig. S2D). *Pira12*, *Lilra6*, *Gapt* and *Pik3r6* represent genes which follow this predominant pattern of PU.1 promoter binding, high expression in control granulocytes and strongly reduced expression after PU.1 deletion (Fig. 2C). In conclusion, this data is strongly supportive of PU.1 as a direct activator of gene expression associated with granulocyte maturation.

**Fig 2.**
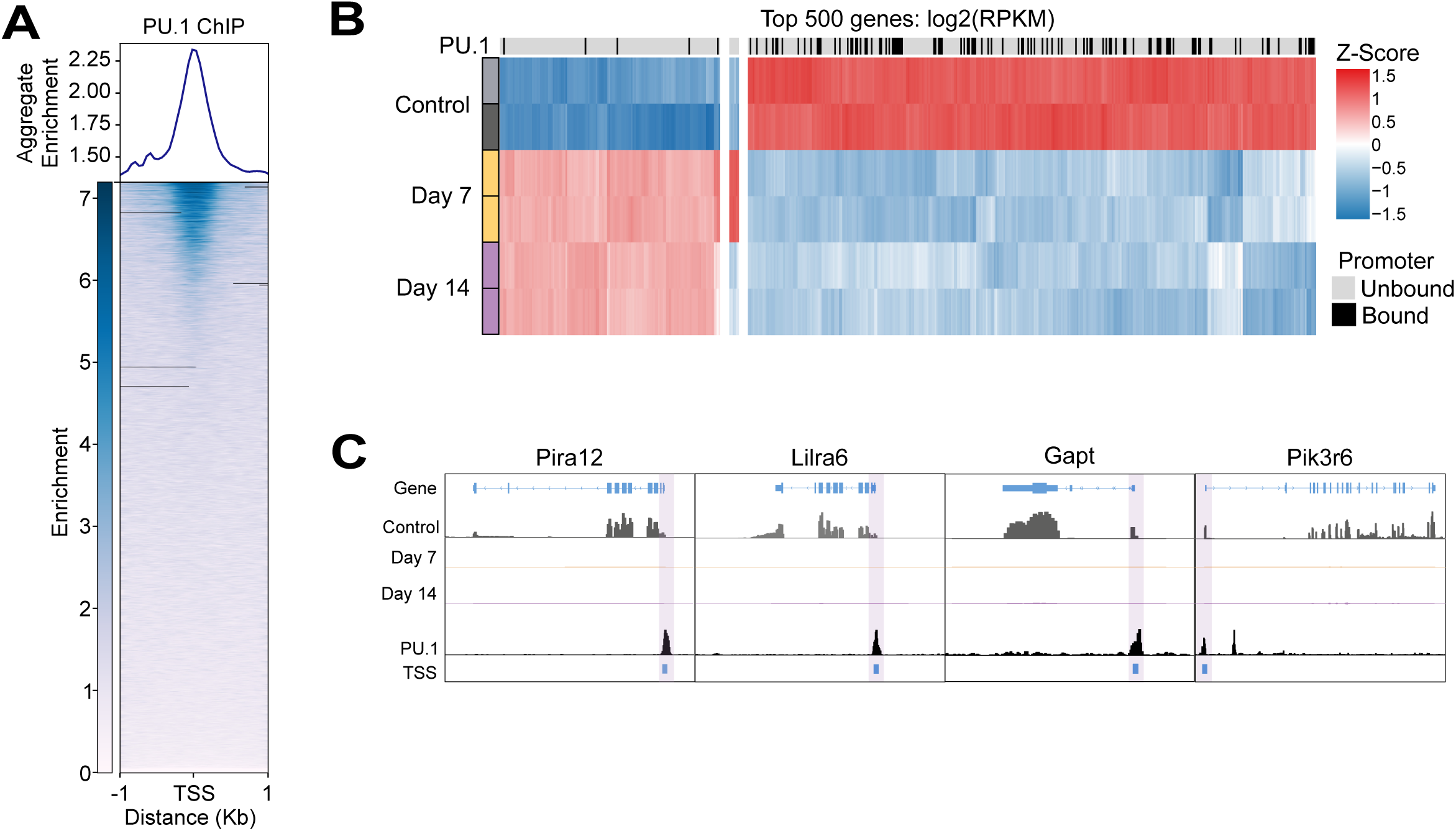
PU.1 binding is associated with active genes in granulocytes. **A**, Heat map showing PU.1 ChIP-seq signal on transcriptional start sites (TSS), ranked by binding intensity. **B**, Heat map displaying RNA-seq expression profiles of top 500 log2(RPKM) genes. Genes with promoter bound by PU.1 are labelled in the heat map. **C,** IGV genome browser tracks showing gene expression and PU.1 ChIP-seq peaks at PU.1 dependant genes. TSS regions are labelled in blue.

### 3D genome architecture is altered by PU.1-inactivation

As PU.1 has been reported to be involved in activating distal enhancers^14, 15^, and more than half of PU.1 binding sites fall outside of promoters (Fig. S2A), we hypothesised that it may play a role in regulating long-range enhancer-promoter interactions during granulocyte maturation. Therefore, we next examined how the 3D chromatin landscape was altered after PU.1 depletion. We performed chromosome conformation capture (*in situ* HiC) and used the diffHiC package to identify over 47189 differential interactions (DIs) PU.1-deficient granulocytes seven days after tamoxifen treatment versus control at 50 kbp resolution (Fig. S3A). Mirroring the DEs from the RNAseq data, there were very few DIs between day 7 and 14 post PU.1 deletion (Fig. S3B).

Next, we directly examined the relationship between PU.1 binding and 3D genome architecture. We focused on the anchors of DIs that were either bound by PU.1 in one, or both anchors in control granulocytes (Fig. 3A). We then asked which direction these DIs trended in PU.1-deficient granulocytes versus control. While the proportion of DIs that were unbound or had binding of PU.1 in one anchor showed an almost equal split in DI direction, DIs that were bound in both anchors were significantly reduced in PU.1-deficient granulocytes (Fig. 3B & Fig. S3C). Moreover, we observed a significant reduction in expression of genes that reside in PU.1-anchored DIs when compared to unbound DIs (Fig. S3D), with examples of these genes shown in Fig. 3C-E. This data suggests that the presence of PU.1, particularly at sites where there is binding at both anchors, is important for the stabilisation of murine granulocyte genome architecture.

**Fig. 3.**
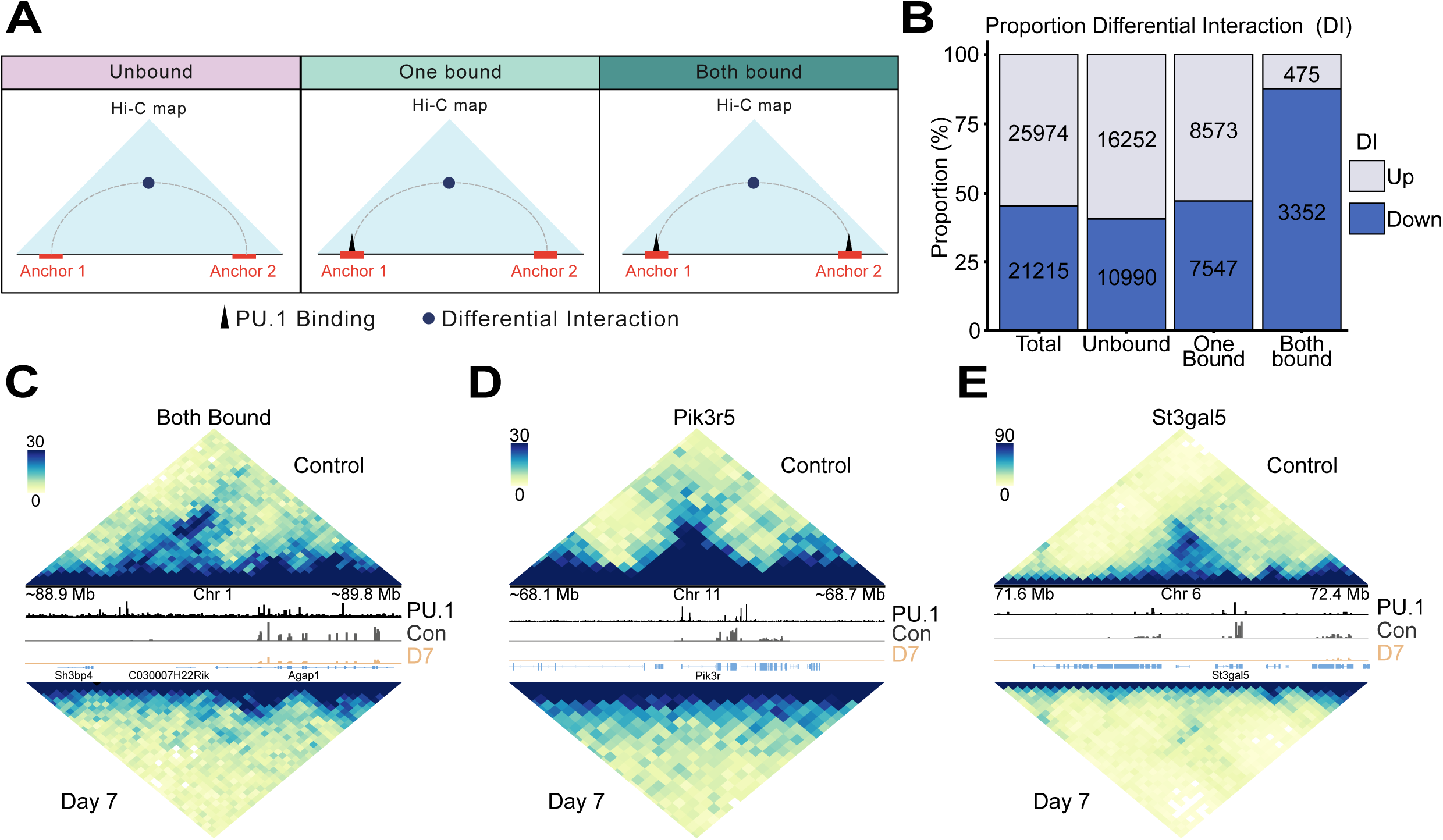
Loss of PU.1 results in reduced genome interactions at PU.1 bound sites. **A**, Schematic depicting classification of PU.1 binding at DI anchors. Unbound are DIs where anchors do not overlap with a PU.1 peak, one bound are DIs with anchors that overlap PU.1 at least once and both bound are DIs with PU.1 overlapping both anchors. **B,** Proportion of DI directionality, colour coded in lilac for down regulated and indigo for up regulated DIs at day 7 post-tamoxifen treatment, with number of DIs shown. **C-E,** *In-situ* Hi-C contact matrix of select regions (20kbp resolution) showing genome architecture. Colour scale indicates number of reads per pixel. RNA-seq and ChIP-seq coverage depicting gene expression and PU.1 binding.

To specifically investigate how chromatin interactivity around gene promoters was affected by loss of PU.1, we examined our HiC data for differential interactions at promoters (DIPs)^25^. We found thousands of gene promoters with altered 3D genome interactions, both increased (>1800) and decreased (>1400) relatively equally (Fig. 4A). As expected, a gain of interactivity was correlated with an increase gene expression and promoters that had reduced interactivity were decreased in gene expression (Fig. 4B). Notably, PU.1 target genes, including *Samhd1*, *Pik3r6* and *Lrrk2* were reduced in both interactivity and expression (Fig. 4B). In line with the role of PU.1 as an activator of gene expression, DIPs that were bound by PU.1 were overwhelmingly reduced in the PU.1-deficient cells (390 down and 62 up, Fig. 4C). To probe this further we inspected the top 500 DIPs and their associated PU.1 binding (Fig. 4D). We observed a clear separation with genes with high interactivity that were reduced in PU.1-deficient granulocytes displaying PU.1 binding whereas genes which gained interactivity displayed little PU.1 binding. Overall, this data suggests that PU.1 binding, and associated 3D genome architecture is linked with the upregulation of genes involved in granulocyte maturation.

**Fig. 4.**
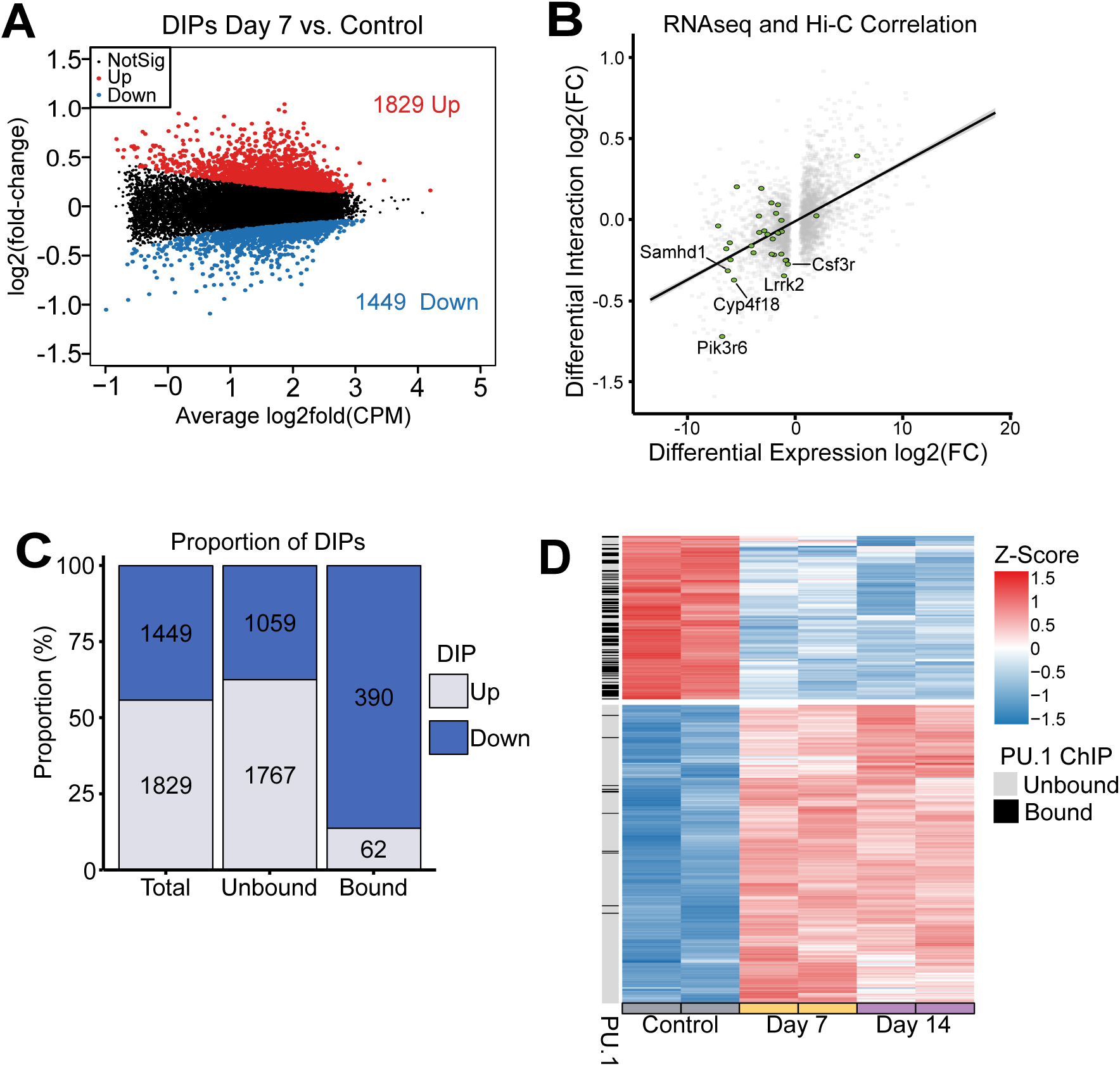
PU.1 deficiency results in reduced genome interactions and gene expression at PU.1 bound sites. **A**, Mean-difference (MD) plot showing differential interactions at promoters (DIPs) between control and PU.1 deficient granulocyte day 7 post-deletion (FDR<0.05). The y-axis shows log2 fold-change and x-axis shows mean abundance log2(CPM). Interactions that are significantly up regulated are highlighted (red) or down regulated are highlighted (blue). **B,** Correlation plot comparing log2(FC) of differential expression and differential interaction filtered significance (FDR <0.05) between control and day 7 post-tamoxifen treatment granulocytes. Top 5 DIPs (determined by highest fold-change) from PU.1 dependent genes are highlighted. **C,** Bar plot showing changes in DIPs between control and day 7 PU.1 depleted granulocytes with and without PU.1 binding. **D,** Heat map displaying top 500 log2(RPKM) DIPs with associated PU.1 binding.

## Discussion

PU.1 is an extensively studied transcription factor with well-established roles in early hematopoiesis and the generation of immune responses^1, 2, 3, 5, 6^. Perturbation of PU.1 levels has been linked to the development of a preleukemic state that precedes acute myeloid leukemia, making understanding how reduction of PU.1 in myelopoiesis extremely important. There is a wealth of knowledge on how PU.1 functions at the molecular level to regulate gene expression. It is a canonical “pioneer” transcription factor which can bind closed chromatin and via interactions with chromatin remodeling factors leads to gene activation^15^. While PU.1 can perform this role at gene promoters, it also plays a prominent role at enhancer regions where it has been implicated in licensing these regions for activation of their target gene(s). Despite this, how PU.1’s involvement in regulating genome architecture more broadly has not been studied. Here we implicate PU.1 as having a role in regulating 3D genome architecture during granulocyte maturation.

We confirm that PU.1 is largely involved in activating gene expression through direct binding at promoters. Analysis of RNAseq data identified ∼6300 DE genes at day 7 after PU.1 depletion (Fig. 1B). As expected, downregulated genes include classic PU.1 targets (Fig. 2C). In the examples shown (*Pira12*, *Lilra6*, *Gapt* and *Pik3r6)* (Fig. 2C), direct binding of PU.1 at the promoter is associated with the expression of these genes. This reaffirms PU.1’s role in mediating granulopoiesis through binding at promoters and activating target genes. In line with previous findings^1, 7^, our results show that disruption of PU.1 leads to a developmental block in granulopoiesis and accumulation of immature granulocytes that maintain cKit expression (Fig. S1A) and fail to upregulate “late neutrophil” genes (Fig. 1C&D).

Intriguingly, despite PU.1’s role as a pioneer factor and reported role in activating transcription through direct targeting, out of the 9001 PU.1 binding sites detected, we found only 987 of these sites to be at promoters of DE genes (Fig. S2C). As PU.1 has previously been reported to act on distal enhancers^14, 15^, we predicted that PU.1 may also be regulating granulopoiesis through long-range enhancer-promoter interactions. To investigate the 3D genome architecture of PU.1 deficient murine granulocytes, we performed *in-situ* HiC and identified over 47189 differential interactions between control and PU.1-deficient granulocytes seven days after tamoxifen treatment (Fig. S3A). Consistent with RNAseq (Fig. S1), changes in genome organisation stabilises after day 7 with relatively few DIs between day 7 and day 14 (Fig. S3B). This suggests that the effects of PU.1 depletion in genome organisation and gene expression is rapid and sustained post day 7 of tamoxifen treatment.

To investigate how PU.1 binding might affect genome architecture in murine granulocytes, we classified differential interactions between control and PU.1-deficient granulocytes (day 7) as either bound at one anchor (one bound), bound at both anchors (both bound) or unbound (Fig. 3A). We observed an equal distribution of increased and decreased interaction between one bound and unbound DIs. Strikingly, DIs that were bound at both anchors were significantly reduced in PU.1 deficient granulocytes (Fig. 3B & Fig. 4C). Suggesting that direct binding of PU.1 at both anchors is required for genome architecture stabilisation at those regions. Furthermore, we observed a trend whereby genes that overlap bound anchors are also significantly reduced in expression (Fig. 4B), as depicted in the examples *Pik3r5* and *St3gal5* (Fig. 1D). Given alteration in PU.1 leads to accumulation of immature granulocytes (Fig. S1A and Fig. 1C), this suggests that PU.1-deficient granulocytes cannot undergo genome organisational changes required for granulocyte maturation. We propose that alterations observed in PU.1 deficient cells are representative of an inability of these cells to establish the genome architecture and transcriptome required for granulocyte maturation. However, given the nature of our experiment design, we cannot exclude the possibility that PU.1 contributes to the maintenance of pre-existing genome architecture required for mature granulocyte identity. Addressing this question would require investigation of early stages of granulocyte differentiation, which has not been conducted in the present study. Nevertheless, our findings demonstrate that genome architecture in murine granulocyte is regulated by PU.1.

In conclusion, this study provides novel insight into the role of PU.1 in granulocyte maturation and identity through the coordinated regulation of gene expression and higher-order chromatin architecture. Given that the loss of PU.1 establishes a preleukemic state, this architectural collapse may represent a critical early event in the pathogenesis of AML. We demonstrate that PU.1 acts as a critical gatekeeper of granulocyte maturation, through stabilisation of mature granulocyte chromatin architecture.

## Methods

### Mice

The *Spi1*^fl^ ^26^ and *Rosa26*-Cre-ER^T2^ ^27^ mice have been previously described. For conditional inactivation of PU.1, *Spi1*^fl/fl^ CreERT2^+^ mice were oral gavaged twice with 5mg Tamoxifen (Sigma, CAS 10540-29-1) in peanut oil, with a day between administrations. Mice were analysed at 7 and 14 days after the first tamoxifen injection. Control mice were untreated *Spi1*^fl/fl^ CreERT2^+^ mice. Experimental mice were used at 6-12 weeks of age and maintained on a C57Bl/6 background. All mice were maintained at The Walter and Eliza Hall Institute Animal Facility under specific-pathogen-free conditions, ambient temperature between 22–24 °C, humidity 40–60%, and light/dark cycle of 12 h/12 h. All aged (200-400 days) females were randomly chosen from the relevant pool. All experiments were approved by The Walter and Eliza Hall Institute Animal Ethics Committee (#2018.002, #2022.002 and #2022.034) and performed under the Australian Code for the care and use of animals for scientific purposes.

### Cell isolation

Washed, single cell homogenate of murine bone marrow was stained with Ly6G-PE (Thermo Fisher, AB_2572720), Ly6C-APC-cy7 (BD, AB_396775), cKit-APC (Thermo Fisher, AB_469430), CD11b-PE-cy7 (Thermo Fisher, AB_469588), TCRb-PerCPcy5.5 (BD, AB_2738629) and CD19-Pacific Blue (Thermo Fisher, AB_10373689). Granulocytes were isolated as single, live, Ly6G^+^, Ly6C^+^, CD19^-^ cells using flow cytometry. CD11b surface expression was also used as a surrogate marker of PU.1 deletion.

### qPCR

RNA was extracted from 100,000 sorted granulocytes from 14-day post-tamoxifen-treated *Spi1* ^fl/fl^ CreER^T2+^ or control mouse bone marrow using chloroform/TRI homogenate/isopropanol precipitation. cDNA was generated using iScript reverse transcription mix (BioRad, #170-8841) following manufacturer’s instructions. *Spi1* (PU.1) and *Hprt* expression levels were then quantified and compared using SensiMix No-Rox kit (Bioline, QT650), following manufacturer’s instructions.

#### Primers

HPRT F – GGGGGCTATAAGTTCTTTGC

HPRT R – TCCAACACTTCGAGAGGTCC

PU.1 F – CTCAGTCACCAGGTTTCCTACA

PU.1 R – AGGTCATCTTCTTGCGGTTG

### RNASeq

RNA was extracted using RNeasy Plus Mini kit (QIAGEN) following manufacturer’s instructions and subsequently quantified using the TapeStation 2200 (Agilent). Libraries were prepared with a TruSeq RNA sample-preparation kit (Illumina) from 500 ng RNA, as per the manufacturer’s instructions. Libraries were then amplified with KAPA HiFi HotStart ReadyMix (Kapa Biosystems) and 200 to 400 bp products were size-selected and cleaned up with AMPure XP magnetic beads (Beckman). Final libraries were quantified with TapeStation 2200 using D1000 ScreenTape for sequencing on the Illumina NextSeq 500 platform to produce 81 bp paired-end reads. 20-50 million read pairs were generated per sample.

### In situ HiC

*In situ* HiC was performed according to Rao, Huntley^73^ and as previously described^25^.

### RNA sequencing and analysis

Reads were aligned to the mm39 genome using Rsubread (v2.14.2) package^28^ and summarised across genes using featureCounts. Genes were identified using version M33 of the GENCODE annotation.

Differential expression (DE) analysis was performed using the R package edgeR^29, 30^ (v4.4.2). Low count genes were filtered using filterByExpr. The remaining counts were converted to log2-CPM and normalised using the trimmed mean of M-values (TMM) method^31^. To identify differentially expressed genes between samples, we used edgeR’s quasi-likelihood pipeline^29, 30^. applied glmTreat with fold-change threshold set to 1.2. P-values were adjusted with multiple testing correction using the Benjamini-Hochberg method^32^ and genes with false discovery rate (FDR) <0.05 were called significant.

Tests for over-representation of gene ontology terms were performed using kegga function from limma^33^ (v3.62.2). Gene set testing on neutrophil early/late signatures, pathways in cancer and PU.1 bound gene promoters were carried out using limma’s fry and visualised using barcode plot function. Neutrophil gene signatures were determined from publicly available single-cell RNAseq dataset^34^. We used the author’s gene-level Spearman correlation values along the neutrophil maturation trajectory to assign genes associated with either early (<-0.1) or late (>0.1) neutrophil categories. PU.1 bound gene promoters were generated from ChIP-seq data and pathways in cancer genes were taken from KEGG database.

Data visualisation used ggplot2^35^ (v3.5.2) and ggrepel^36^ (v0.9.6) to generate scatter and barplots. Colour palettes were generated from viridis^37^ (v0.6.5) and RColorBrewer^38^ (v1.1-3).

### PU.1 ChIP sequencing, data processing and analysis

ChIP samples were prepared on C57Bl/6 granulocytes according to the standard Millipore/Upstate protocol and using the polyclonal anti-PU.1 IgG (T-21 X: sc-352 X) from Santa Cruz. Libraries were prepared and sequenced using the Illumina TruSeq workflow.

ChIP-seq reads were aligned to the mm39 genome using Bowtie2 (v2.4.4), technical replicate BAM files were sorted, indexed and merged using SAMtools^39^ (v1.19.2). Duplicates were marked and removed using Picard-Tools^40^ (v2.6.11) with MarkDuplicates. Peaks were called with MACS2^41^ (v2.2.7.2) on default parameters. To profile PU.1 occupancy around promoters, coverage bigwig files were generated with bamCoverage from Deeptools^42^ (v3.5.1). Transcriptional start sites (TSSs) were defined as regions 1bp upstream and downstream of a gene body using the promoters function from IRanges^43^ (v2.34.1). A TSS anchored matrix was then computed using computeMatrix with paramerters set to 50kb bin size and 1kb flanking regions. Heat map was generated using plotHeatmap from Deeptools (v3.5.1). Genomic ranges for DE genes and PU.1 peaks were constructed using the Granges function in GenomicRanges (v1.52.0) R package^43^. DE gene promoters were determined using the promoters function from IRanges^43^ (v2.34.1) with default parameters (2kb upstream and 200bp downstream TSS). Intersection between DE gene promoter and PU.1 peaks were performed using the find_overlaps function from plyranges^44^ (v1.20.0). ENTREZ IDs were converted from gene symbols using synGO^45^.

### HiC processing

In situ HiC was performed on fixed cells as previously described^46^. Independent biological duplicates were prepared for each condition and sequenced to a depth of around 200 million reads (Supplemental Table 1).

HiC libraries were aligned to the mouse reference genome (mm39) using pre_split_map.py script from the diffHic (v1.38.0) package^47^, which splits reads at ligation junctions and aligns the fragment with bowtie2 (v2.4.4). Technical replicates were merged using SAMtools (v1.19.2) and BAM files were processed with Picard-Tools, mate pair information was synced using FixMateInformation, duplicate reads were marked with MarkDuplicates^40^. The resulting bam files were then sorted by read name using SAMtools (v1.19.2). Read pair processing were analysed following the diffHic pipeline^47^ as previously described^48^. A param object was generated using the BSgenome.Mmusculus.UCSC.mm39 package (v.1.4.3), the genome was digested with Mbo1 recognition motif using the cutGenome function. Using preparePairs function, read pairs were mapped to specific Mbo1 restriction fragment and valid reads were identified. A read pair was discarded if the read was unmapped, marked as duplicate, or had a mapping quality score below 10. Additional filtering removed artifacts such as self-circling and dangling ends, defined as inward-facing or out-ward facing reads on the same chromosome separated by <1kb. To visualise HiC data, biological replicates were merged using the mergePairs function in diffHic and converted into hic or cool format using Juicer^49^ (v1.22.01). Contact matrices were generated using the plot_hic_region function in plotgardener (v.1.12.0) at 30kb resolution.

### HiC differential analysis

Differential interactions (DIs) between samples were detected using the diffHic package (v1.38.0) as previously described^48^. Read pairs were counted into 20kb bin size using the squareCounts function from diffHic. Interacting bin pairs were filtered on interaction intensities 5-fold greater than background ligation frequency using filterDirect. Background ligation frequency was estimated from inter-chromosomal bin pairs determined from a 2Mbp bin pair count matrix. Bin pair counts were normalised between libraries using loess-based approach with normOffset. To test for DIs, we implemented the quasi-likelihood framework in edgeR^29, 30^. Significant DIs were assessed using Benjamini–Hochberg corrected FDR threshold set to <0.05. DIs were classified as one bound, both bound or unbound using the R package InteractionSet^50^ (v1.28.1). An anchor was considered bound if it overlapped with a MACS2 called PU.1 peak using the find_overlaps function with default parameters. DIs with one bound anchor were labelled one bound; those with both anchors bound were both bound and those with no bound anchors were unbound. DE genes at DI anchors were also identified using the findOverlaps from InteractionSet^50^ (v1.28.1).

### Differential interacting promoter analysis

To identify differential interacting promoter (DIP), we used the diffHic package (v1.38.0) as described previously^25, 48^ with alteration in certain parameters. Promoter regions were defined as 9kb upstream and 1kb downstream of gene TSS using the mm39 refseq annotation from the Rsubread (v2.14.2) package^28^. Promoter-anchored interactions were counted using the connectCounts function from diffHic with bin size set to 10kb. Low abundance interactions that did not exceed the fitted loess trend were filtered out using parameters previously described^48^. Additionally self-ligation events (interactions along the diagonal) or excessively large anchors >15kb were also filtered out. Interactions for each promoter were aggregated by gene name to generate a counts matrix representing the summed interactions per gene promoter. Gene promoters with low counts were filtered out using edgeR’s filterByExpr with the following parameters: min.count=100, min.total.count =150. Only promoters for protein encoding genes were retained for analysis. Genes annotated as rRNA, ncRNA, mitochondrial, Riken and olfactory receptor genes were excluded from downstream analysis. Counts were normalised using loess method and DIPs were detected using the edgeR quasi-likelihood framework.

### Plotting

MA and MDS plots were visualised using limma (v3.62.2). Heat maps were generated using pheatmap^51^ (v1.0.12).). Bar plots and box plots were generated using ggplot2^35^ (v3.5.2) and colour palette from viridis^37^ (v0.6.5) and RColorBrewer^38^ (v1.1-3).

---

**Fig. S1.**
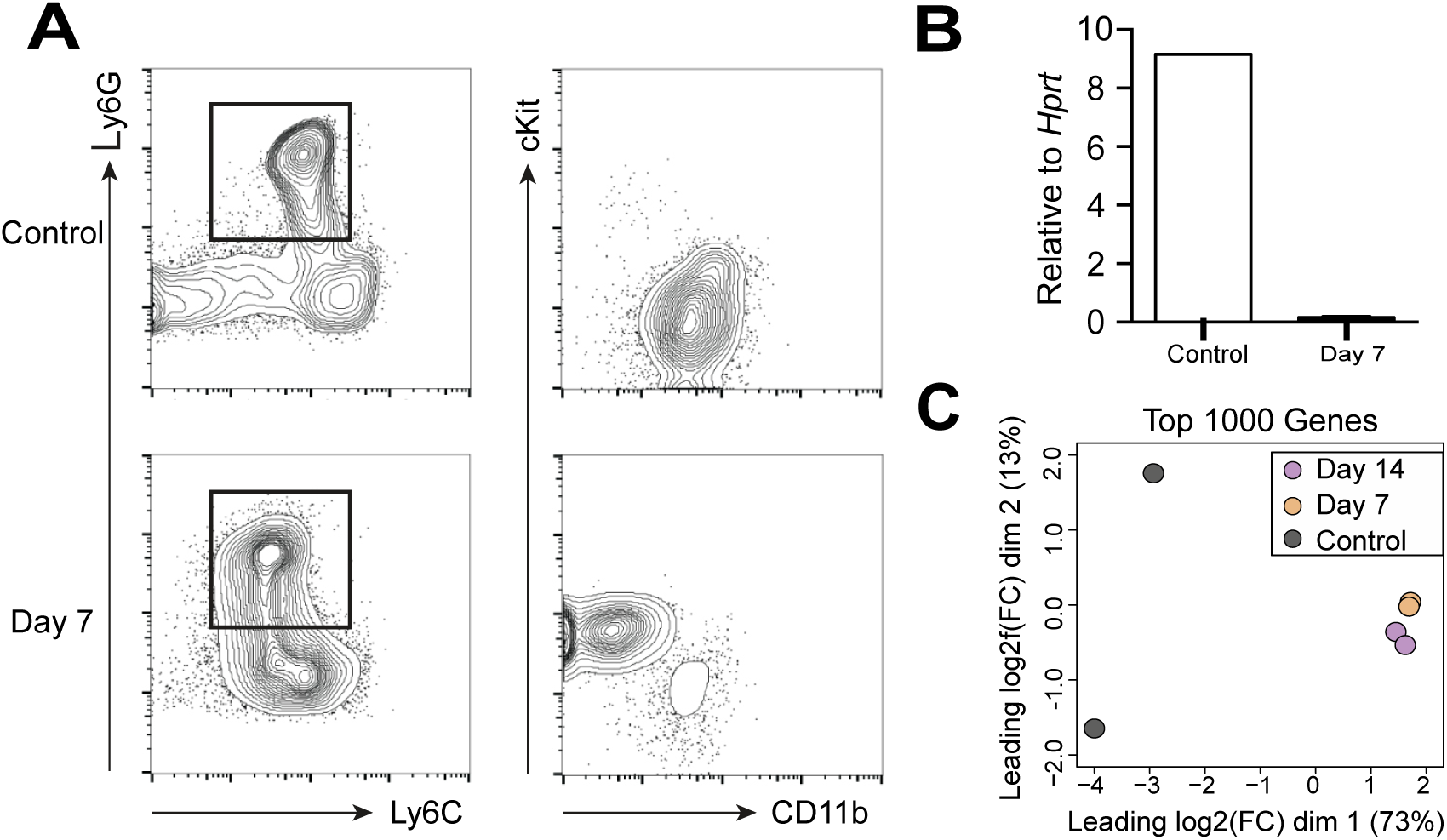
Transcriptome wide alterations in PU.1-deficient granulocytes. **A**, Flow cytometry of cells from *Spi1*fl/fl CreERT2+ mice either untreated (control) or days 7 post-tamoxifen treatment (Day 7). Granulocytes were isolated using the following gating strategy: CD19-Ly6G+Ly6C+. cKit and CD11b expression was also assessed. **B**, qPCR results confirming PU.1 KO in granulocytes 7 days post tamoxifen treatment, *Spi1* expression was normalised relative to *Hprt* expression. **C**, Multidimensional-scaling plot visualising the gene expression relationships in mouse granulocytes following PU.1 deletion at days 7 and 14 post-tamoxifen treatment. Distances on the plot correspond to leading log_2_fold-change (FC), computed as the root-mean-square average of the top 1000 largest log_2_FC sample pairs.

**Fig S2.**
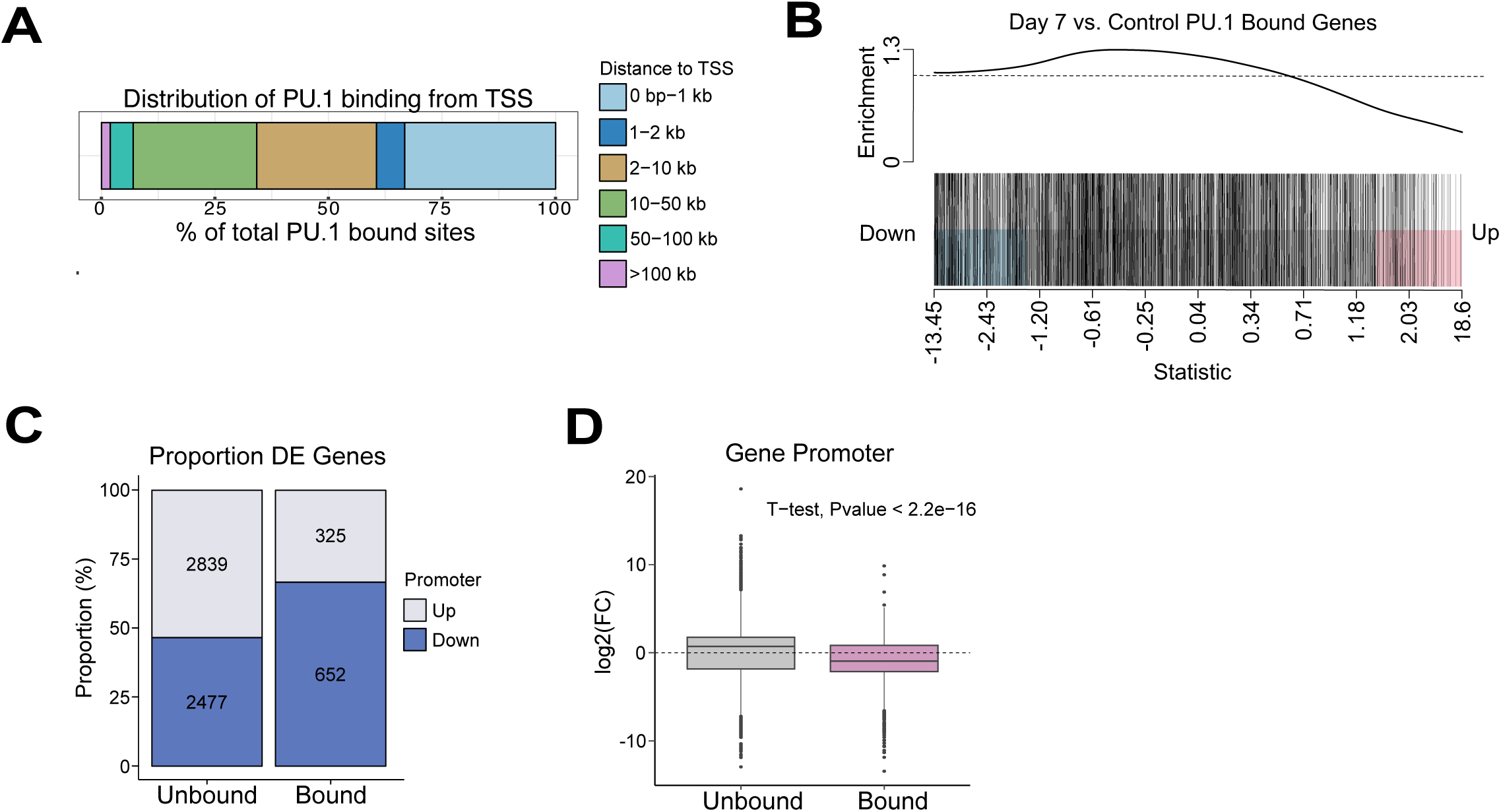
PU.1 binding is associated with active genes in granulocytes. **A**, Distribution of PU.1 binding sites from the TSS represented as % of binding sites. **B,** Barcode plot representing enrichment of genes with PU.1 bound at promoter. Each straight bar corresponds to a specific gene (red for up regulation and blue for down regulation) and the line above show relative enrichment. Statistics represent logFC ranking of genes. **C**, Proportion of DE gene directionality, colour coded in blue for down regulated (Log_2_FC < 0) and red for up regulated (log_2_FC < 0) genes at Day 7 post-tamoxifen treatment. DE gene list was split into PU.1 bound and unbound gene promoters. **D**, Box plot depicting DE gene log_2_FC (Day 7 vs untreated) at PU.1 bound and unbound promoters. Statistical significance was determined using an unpaired two sample t-test.

**Fig. S3.**
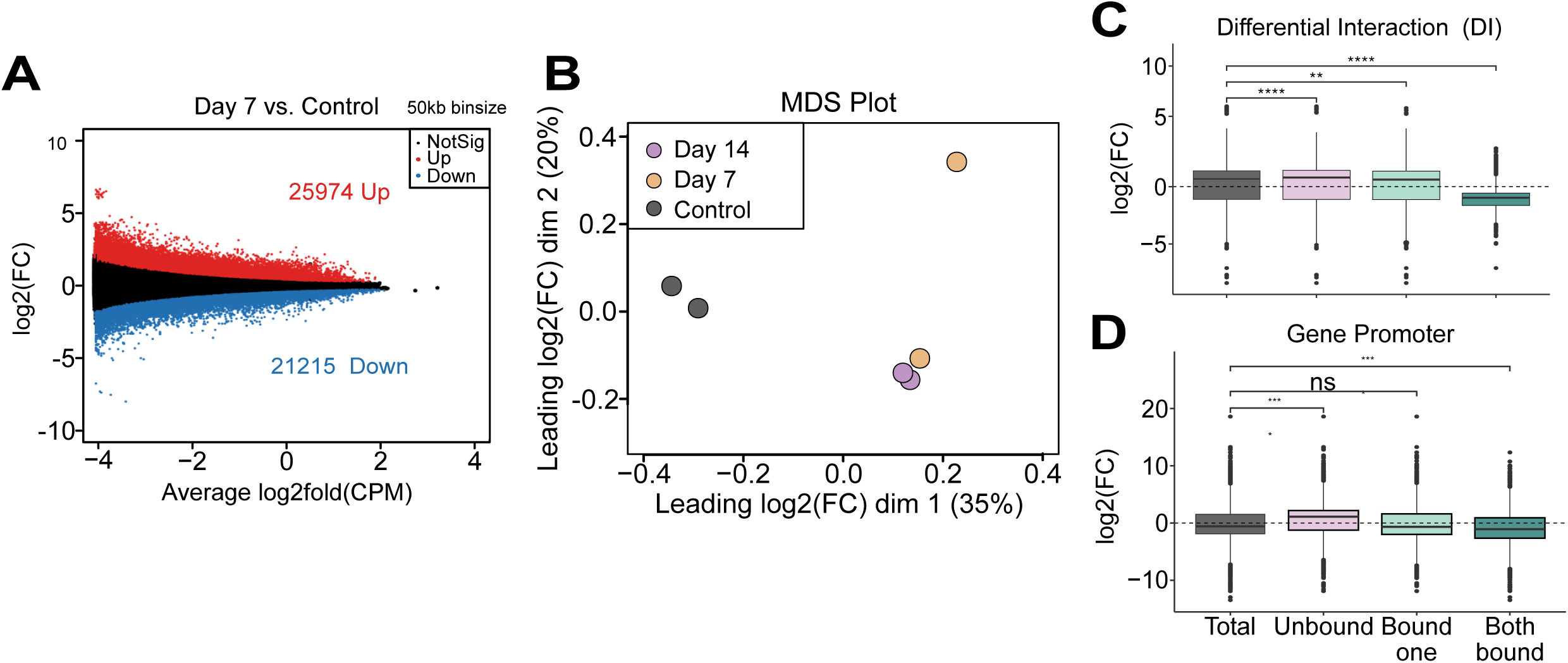
PU.1 deficiency results in altered genome interactions and gene expression at PU.1 bound sites. **A**, Mean-difference (MD) plot showing differential interaction between wild type murine and PU.1 deficient granulocyte day 7 post-deletion (FDR<0.05). The y-axis shows log2 fold-change and x-axis shows mean abundance log2(CPM). Interactions that are significantly up regulated are highlighted (red) or down regulated are highlighted (blue). **B,** Multidimensional-scaling (MDS) plot of in-situ Hi-C data of murine granulocytes following PU.1 deletion at day 7 and 14 post-tamoxifen treatment. **C,** Box plot showing log2(FC) between control and day 7 post-tamoxifen treatment at unbound, one bound and both bound DIs. **D,** Box plot depicting log2(FC) of differential expressed genes between control and day 7 post-tamoxifen treatment that overlap anchors of DIs that are unbound, one bound and both bound.

